# Mitochondrial impairment and rescue in riboflavin responsive neuropathy

**DOI:** 10.1101/110890

**Authors:** Andreea Manole, Zane Jaunmuktane, Iain Hargreaves, Amelie Pandraud, Vincenzo Salpietro, Simon Pope, Marthe H. R. Ludtmann, Alejandro Horga, Renata S. Scalco, Abi Li, Balasubramaniem Ashokkumar, Charles M. Lourenço, Simon Heales, Rita Horvath, Patrick F. Chinnery, Camilo Toro, Andrew B. Singleton, Thomas S. Jacques, Andrey Y. Abramov, Francesco Muntoni, Michael G. Hanna, Mary M. Reilly, Tamas Revesz, Dimitri M. Kullmann, James E.C. Jepson, Henry Houlden

**Affiliations:** Department of Molecular Neuroscience and Neurogenetics Laboratory; MRC Centre for Neuromuscular Diseases; Division of Neuropathology and Department of Neurodegenerative Disease; Neurometabolic Unit, National Hospital for Neurology and Neurosurgery, London, WC1N 3BG,UK; Reta Lila Weston Institute of Neurological Studies and Queen Square Brain Bank for Neurological Disorders, Queen Square, London WC1N 3BG, UK; School of Biotechnology, Madurai Kamaraj University, Madurai 625021, India; Departamento de Neurociências e Ciências do Comportamento, Faculdade de Medicina de Ribeirão Preto, Universidade de São Paulo (USP), Ribeirão Preto, SP, Brazil; Chemical Pathology, Great Ormond Street Children's Hospital, London, UK; John Walton Muscular Dystrophy Research Centre, Institute of Genetic Medicine, Newcastle University, Central Parkway, Newcastle upon Tyne, UK; Department of Clinical Neurosciences, School of Clinical Medicine, University of Cambridge, Cambridge CB2 0QQ, UK & MRC Mitochondrial Biology Unit, Cambridge Biomedical Campus, Cambridge CB2 0QQ; NIH Undiagnosed Diseases Program, Common Fund, Office of the Director, Bethesda, MD, USA; Laboratory of Neurogenetics, NIA/NIH, Bethesda, MD, USA; Developmental Biology and Cancer Programme, UCL Great Ormond Street Institute of Child Health, 30 Guilford Street, London WC1N 1EH, UK; The Dubowitz Neuromuscular Centre, UCL Great Ormond Street Institute of Child Health, 30 Guildford Street, London, WC1N 1EH, UK; Department of Clinical and Experimental Epilepsy, UCL Institute of Neurology, Queen Square, London, WC1N 3BG, UK

**Keywords:** Brown-Vialetto-Van Laere syndrome, *SLC52A2*, *SLC52A3*, riboflavin

## Abstract

Brown-Vialetto-Van Laere syndrome (BVVLS) represents a phenotypic spectrum of motor, sensory, and cranial nerve neuropathy, often with ataxia, optic atrophy and respiratory problems leading to ventilator-dependence. Loss-of-function mutations in two riboflavin transporter (RFVT) genes, *SLC52A2* and *SLC52A3,* have recently been linked to BVVLS. However, the genetic frequency, neuropathology and downstream consequences of RFVT mutations have previously been undefined. By screening a large cohort of 132 patients with early-onset severe sensory, motor and cranial nerve neuropathy we confirmed the strong genetic link between RFVT mutations and BVVLS, identifying twenty-two pathogenic mutations in *SLC52A2* and *SLC52A3,* fourteen of which were novel. Brain and spinal cord neuropathological examination of two cases with *SLC52A3* mutations showed classical symmetrical brainstem lesions resembling pathology seen in mitochondrial disease, including severe neuronal loss in the lower cranial nerve nuclei, anterior horns and corresponding nerves, atrophy of the spinothalamic and spinocerebellar tracts and posterior column-medial lemniscus pathways. Mitochondrial dysfunction has previously been implicated in an array of neurodegenerative disorders. Since riboflavin metabolites are critical components of the mitochondrial electron transport chain (ETC), we hypothesized that reduced riboflavin transport would result in impaired mitochondrial activity, and confirmed this using *in vitro* and *in vivo* models. ETC complex I and complex II activity were decreased in *SLC52A2* patient fibroblasts, while global knockdown of the single *Drosophila* RFVT homologue revealed reduced levels of riboflavin, downstream metabolites, and ETC complex I activity. RFVT knockdown in *Drosophila* also resulted in severely impaired locomotor activity and reduced lifespan, mirroring patient pathology, and these phenotypes could be partially rescued using a novel esterified derivative of riboflavin. Our findings indicate mitochondrial dysfunction as a downstream consequence of RFVT gene defects in BVVLS and validate riboflavin esters as a potential therapeutic strategy.

## Introduction

Brown-Vialetto-Van Laere syndrome (BVVLS) is an autosomal recessive neurological disorder first described by Brown in 1894 and later by Vialetto and Van Laere (Brown, 1894; Van Laere, 1966; Vialetto, 1936). Affected patients mostly present with neuropathy, bilateral sensorineural deafness, bulbar palsy and respiratory compromise. Other cranial nerve palsies, optic atrophy, upper and lower motor neuron involvement and ataxia can occur particularly as disease progresses, mimicking conditions such as amyotrophic lateral sclerosis (ALS), Madras motor neuron disease and Nathalie syndrome (Anand et al., 2012; Manole et al., 2014). Deafness is the most common sign of this condition, with most affected individuals exhibiting hearing loss during the disease course. The time between the onset of deafness and the development of other manifestations varies but is usually in early childhood (Manole and Houlden, 2015).

Previous work has revealed strong links between mutations in two genes *(SLC52A2* and *SLC52A3)* and BVVLS, both of which encode riboflavin transporters (RFVTs) (Foley et al., 2014; Green et al., 2010; Johnson et al., 2012). The role of another RFVT-encoding gene, *SLC52A1,* in BVVLS pathogenicity is still uncertain, as it was found to be defective in only one case (Ho et al., 2011). *SLC52A2* and *SLC52A3* mutations include missense, nonsense, frame-shift, and splice-site alterations, but uniformly result in loss-of-function through reduced RFVT expression and/or riboflavin uptake (Foley et al., 2014; Intoh et al., 2016; Udhayabanu et al., 2016).

Riboflavin (7,8-dimethyl-10-ribityl-isoalloxazine) is a water-soluble compound and acts as a precursor for flavin mononucleotide (FMN) and flavin adenine dinucleotide (FAD). Both FMN and FAD function in biological redox reactions such as in the mitochondrial electron transport chain (ETC) (Powers, 2003). Since riboflavin cannot be synthesised by mammals *de novo,* RFVTs are indispensable for normal cellular metabolism, suggesting that reduced intracellular riboflavin is a critical pathological mediator of BVVLS. Indeed, although infants with early-onset RFVT deficiency rapidly become ventilator-dependent and usually die in the first year of life, treatment with high-dose riboflavin supplementation partially ameliorates the progression of this neurodegenerative condition, particularly if initiated soon after the onset of symptoms (Foley et al., 2014).

Motor neurons are thought to be uniquely susceptible to impaired energy metabolism because of their high metabolic rate and axonal length (Cozzolino and Carri, 2012). Furthermore, mitochondrial perturbations causing alteration of the ETC and increased oxidative stress are known to be involved in the pathomechanisms of neurodegeneration in motor neuron diseases such as ALS or spinal muscular atrophy (Bartolome et al., 2013; Cozzolino and Carri, 2012; Johnson et al., 2010). Given the critical role of riboflavin in the generation of substrates used for the ETC, we hypothesized that reduced riboflavin transport results in impaired ETC, which may in turn contribute to neurodegeneration.

Here, we expand the clinic-genetic spectrum of riboflavin transporter genes and then perform a series of experiments to identify the underlying effects of loss of RFVT function on neuronal integrity and mitochondrial function. We review clinical case histories and undertake pathological evaluation of brain and spinal cord of two patients with confirmed *SLC52A3* mutations who presented either in infancy or in later childhood. Finally, we investigate the *in vitro* cellular effects of *SLC52A2* mutations on metabolism and ETC function and also the *in vivo* consequences of loss of the *SLC52A3* homologue in *Drosophila,* and test whether these can be mitigated by supplementation with a riboflavin derivative.

## Methods

### Study Subjects

Patients were enrolled with informed consent from the patient and/or their parental guardian. DNA was collected from a total of 132 suspected cases (probands and their relatives) presenting with cranial neuropathies and sensorimotor neuropathy with or without respiratory insufficiency. Patients’ DNA samples were collected at medical centres in England (including from patients originating from Pakistan, India, Saudi Arabia, Kuwait, Iran and Turkey) as well as from medical centres in Wales, Scotland, Northern Ireland, Ireland, France, Belgium, the Netherlands, Greece, Malta, Russia, Lebanon, Iceland, Australia and the United States, following the announcement of an on-going molecular study at the UCL Institute of Neurology, University College London (National Hospital for Neurology & Neurosurgery, Queen Square, London) of patients presenting with this phenotype. This study was ethically approved by the UCL/University College London Hospital Joint Research Office (99/N103), and written informed consent to perform a skin biopsy and fibroblasts was obtained as appropriate.

### PCR and Sanger sequencing

Primer sequences, PCR and Sanger sequencing conditions for *SLC52A1, SLC52A2* and *SLC52A3* were as in (Foley et al., 2014). Segregation of pathogenic variants was also assessed. Where one heterozygous mutation was identified, deletions in the other allele were investigated by array CGH but no deletions or insertions were identified. Mutation positions are based on NCBI reference sequences for complementary DNA. *SLC52A2* mutation positions are based on sequences NM_024531.4 for the nucleotide sequence and NP_078807.1 for the protein sequence. *SLC52A3* mutation positions are based on sequences NM_033409.3 for the nucleotide sequence, and NP_212134.3 for the protein sequence.

### Neuropathological evaluation

Formalin-fixed, paraffin embedded brain and spinal cord tissue were available from two patients (AM2 and AM4). The paraffin sections were cut at 4 μm, mounted on glass slides and stained with routine H&E and Luxol fast blue/cresyl violet histochemical stains. Sections were examined by immunohistochemistry with the following antibodies: glial fibrillary acid protein (GFAP) (polyclonal, 1:2500, Dako), phosphorylated neurofilaments (clone SMI31, 1:5000, Sternberg), myelin basic protein (clone SMI94, 1:2000, Sternberg), ubiquitin (polyclonal, 1:1200, Dako), p62 (3/P62LCK Ligand, 1:100, BD Transduction), α-synuclein (clone KM51, 1:50, Leica/Novocastra), amyloid-β (clone 6F3D, 1:100, Dako), hyperphosphorlyated tau (clone AT8, 1:1200, INNOGENETICS), TDP43 (clone 2E2-D3, 1:3000, Abova), CD68 (clone PG-M1, 1:100, Dako), CD3 (LN10, 1:100, Leica/Novocastra), CD20 (clone 7D1, 1:200, Dako). Immunohistochemistry was carried out either manually or on a BondMax autostainer (Leica Microsystems) using 3,3-diamunobenzidine as chromogen. Negative controls were treated identically except that the primary antibody was omitted. Appropriate positive controls were used for all immunohistochemical studies.

### *SLC52A2* patient fibroblasts

#### Generation of human fibroblast cultures and cell culture

Skin fibroblasts of BVVLS patients with *SLC52A2* mutations were generated at the Medical Research Council (MRC) Centre for Neuromuscular Diseases Biobank, Dubowitz Centre, UCL Institute of Child Health (ICH) by Dr Diana Johnson, or sent in culture by collaborators. Three age-matched controls were obtained from the ICH Biobank: Control 1 (C1) (age at biopsy: 14 years; female); Control 2 (C2) (age at biopsy: nine years; male); Control 3 (C3) (age at biopsy: five years; male). Five BVVLS patients’ fibroblast lines were available for this study. Patients carried the following mutations: E1:p.[Gly306Arg];[Gly306Arg];E2:p.[Trp31Ser];[Leu312Pro];E3:p.[Gln234Stop];[Ala420Thr]; E4:p.[Gly306Arg];[Leu339Pro]; I1: p.[Tyr305Cys]; [Gly306Arg]. All cells were maintained at 37°C and 5% CO_2_ under humidified conditions and cultured in high glucose Dulbecco’s modified Eagle medium (Invitrogen) supplemented with 10% foetal bovine serum (Biowest). All fibroblast lines were grown for four days in modified DMEM containing physiological concentrations (12.6 nM) of riboflavin, followed by four days in modified DMEM, which was either riboflavin-supplemented (300.6 nM) or contained a low riboflavin concentration (3.1 nM).

#### Determination of Flavin status in fibroblasts

Riboflavin, FMN and FAD content were measured in neutralized perchloric extracts by means of High Performance Liquid Chromatography (HPLC), as previously described (Giancaspero et al., 2009). Quantitative determination of riboflavin, FMN, and FAD was carried out using the Whole Blood Chromsystems vitamins B_1_/B_2_ kit (Chromsystems, Germany) as per the manufacturer’s protocol. The Bio-Rad DC protein assay (Bio-Rad Laboratories, USA) was used to normalise for protein concentration.

#### Assessment of ETC complex I, II, and citrate synthase activities in fibroblasts

All enzyme activities were determined at 30^o^C. Prior to analysis all samples were subjected to three cycles of freeze/thawing to lyse membranes. Enzymatic activities were determined using an Uvikon 940 spectrophotometer (Kontron Instruments Ltd, Watford, UK).

Complex I activity was measured according to the method of (Ragan, 1987), which involved monitoring the oxidation of NADH at 340 nm. Complex II assay was measured according to the method of (Birch-Machin et al., 1994), which monitored the succinate-dependent 2-thenoyltriflouroacetone sensitive reduction of 2,6-dichlorophenolindophenol at 600 nm. The activity of citrate synthase was measured by the formation of the anion of thionitrobenzoate from 5,5'-dithiobis(2-nitrobenzoate) and CoA at 412 nm (Shephard, 1969). This provided an estimate of mitochondrial content and was therefore used to normalise complex I and II activities for mitochondrial enrichment (Hargreaves et al., 1999).

#### *Drosophila* stocks and culture conditions

Fly strains were obtained from the Bloomington *Drosophila* Stock Center (Indiana, USA) and Vienna *Drosophila* Resource Center (Austria). All transgenic insertions used in this study were outcrossed at least 5 times into an isogenic (iso31) background. These were: *cg11576* UAS-RNAi ((VDRC 7578)

(Dietzl et al., 2007) and HMC04813 (Perkins et al., 2015)), and *daughterless*-GAL4 (Bloomington stock 55850). Flies were reared at 25°C on standard fly food consisting of corn meal, yeast, sucrose, glucose, wheat germ, soya flour, nipagin and propionic acid. For experiments with compound supplementation, 0.1 mg/ml riboflavin (Sigma) or 0.1 mg/ml riboflavin-5'-lauric acid (RLAM) was added to the food. Flies were kept under 12 h light: 12 h dark cycles, defined as LD conditions.

#### Semi-quantitative RT-PCR for whole flies

Total RNA was extracted from flies using TRIzol, as per manufacturer’s instructions. The concentration and purity of RNA was determined spectrophotometrically. 1 μg of RNA was reverse transcribed to first strand cDNA by using random primers and Moloney murine leukemia virus reverse transcriptase (Promega, Madison, WI, USA). Primers used for *cg11576 (drift)* were designed so that they span exon-exon boundaries, Forward– 5’ CCAGATGCTCCTCTCTCGA 3’ Reverse– 5’ AGTACACAGTCGCCACTCTC 3’. Primers used for *Drosophila rp49* were: Forward– 5’ CCCAACCTGCTTCAAGATGAC 3’, Reverse – 5’ CCGTTGGGGTTGGTGAGG 3’. GoTaq^®^ Green Master Mix was used and PCR reactions were performed with the following protocol: 95°C-2 min, (95°C-30 s, 60°C-30 s, 73°C-1 min) for 35 cycles, 73°C-5 min, and 4°C-hold. Two exponential curves representing the product formation was made for both primer pairs and cycles 26 and 29 were chosen for *rp49* gene and *drift* respectively so that amplification rates were in the linear range for semi-quantitative comparisons.

#### Quantification of Flavins, ETC complex I, II, II/III redox activities and citrate synthase assay in *Drosophila*

Riboflavin, FMN and FAD content, and ETC complex I, II redox activity and citrate synthase activities were measured in flies (approximately 10/genotype) as described above for patient fibroblasts. Prior to analysis all samples were subjected to three cycles of freeze/thawing to lyse membranes. A Lowry assay was used to normalize for protein concentration (Frolund et al., 1995). Complex II/III activity was determined at 30°C using the method of (King, 1967) which followed the succinate-dependent antimycin A sensitive reduction of cytochrome *c* at 550 nm.

#### Larval behaviour

Larval locomotion was tested by placing individual third instar larvae in the center of petri dishes (8.5 cm diameter, 1.4 cm height) coated with 10 ml of 4% agar. On average, 30 larvae were tested per strain. The number of grid squares (1 cm) entered per min by the larva was analysed using Kruskal-Wallis tests and subsequently Dunn’s post-hoc tests.

#### Adult behaviour

Adult flies were kept as groups of males and females in a 12h:12h light-dark (LD) cycle at 25°C for 1 day prior to testing. Single virgin females (approx. 1 day old) were loaded into glass tubes (with 2% agar and 4% sucrose food) and monitored using the *Drosophila* Activity Monitoring System (Trikinetics) in LD at 25°C with an approximate intensity of 700-1000 Lux during the L condition. For experiments involving supplementation, riboflavin or RLAM was added to the above food. Fly activities were deduced from the number of times flies broke beams of infrared light passing through the middle of the tube. Locomotor activity was recorded in 30 min bins and an analysis was performed on the second day after loading. Data were pooled from at least two independent experiments. The relative locomotor activities per 30 min bin for individual flies were averaged for each genotype and also the average locomotor activity per day was calculated. Locomotion graphs were generated using GraphPad Prism 6 and Microsoft Excel.

#### Life span

Adult female flies were collected from eclosion and transferred to fresh food tubes (10 flies/tube) with or without RLAM supplementation. Each day, death events were scored and viable flies were transferred to fresh tubes. Survival proportions were plotted as percentage of live flies against days. Approximately 100 flies were tested for each genotype.

#### Other statistics

Statistics were performed using GraphPad Prism 6. The significance between the variables was shown based on the p-value obtained (ns indicates p > 0.05, * indicates p < 0.05, ** indicates p < 0.005, *** indicates p < 0.0005). Data are presented as box plots illustrating 80% of the data distribution, together with the median and 10^th^, 25^th^, 75^th^ and 90^th^ percentiles.

## Results

### Clinical and genetic analysis of BVVLS patients

We screened 132 patients with phenotypes suggestive of BVVLS and identified twenty patients (15%) with RFVT mutations. Genetic and clinical details of the twenty individuals carrying mutations in *SLC52A2* or *SLC52A3* are summarised in Table 1. Thirteen of the probands were males. Age of symptom onset was available for 15 patients (mean: 8.2 years; range: 7 months - 21 years). First symptoms usually occurred during childhood (11 patients) and less frequently during teenage years (four patients) or adulthood (one patient), and were mostly secondary to cranial nerve involvement (Table 1). Only one patient presented with symptoms not related to cranial nerve involvement (limb weakness). Sensorineural hearing loss was both a common presenting symptom (eight patients) and a common clinical feature during follow-up (17 patients). Other frequent clinical features include optic atrophy (14 patients), weakness of facial and bulbar muscles (13 patients) and sensorimotor peripheral axonal neuropathy (16 patients). Limb weakness was more severe in the upper than in the lower limbs in nine patients. Ten patients developed some degree of respiratory involvement with three requiring assisted ventilation, and ten patients had dysphagia and/or chewing difficulty, six of them requiring nasogastric tube or gastrostomy feeding.

**Table 1.**
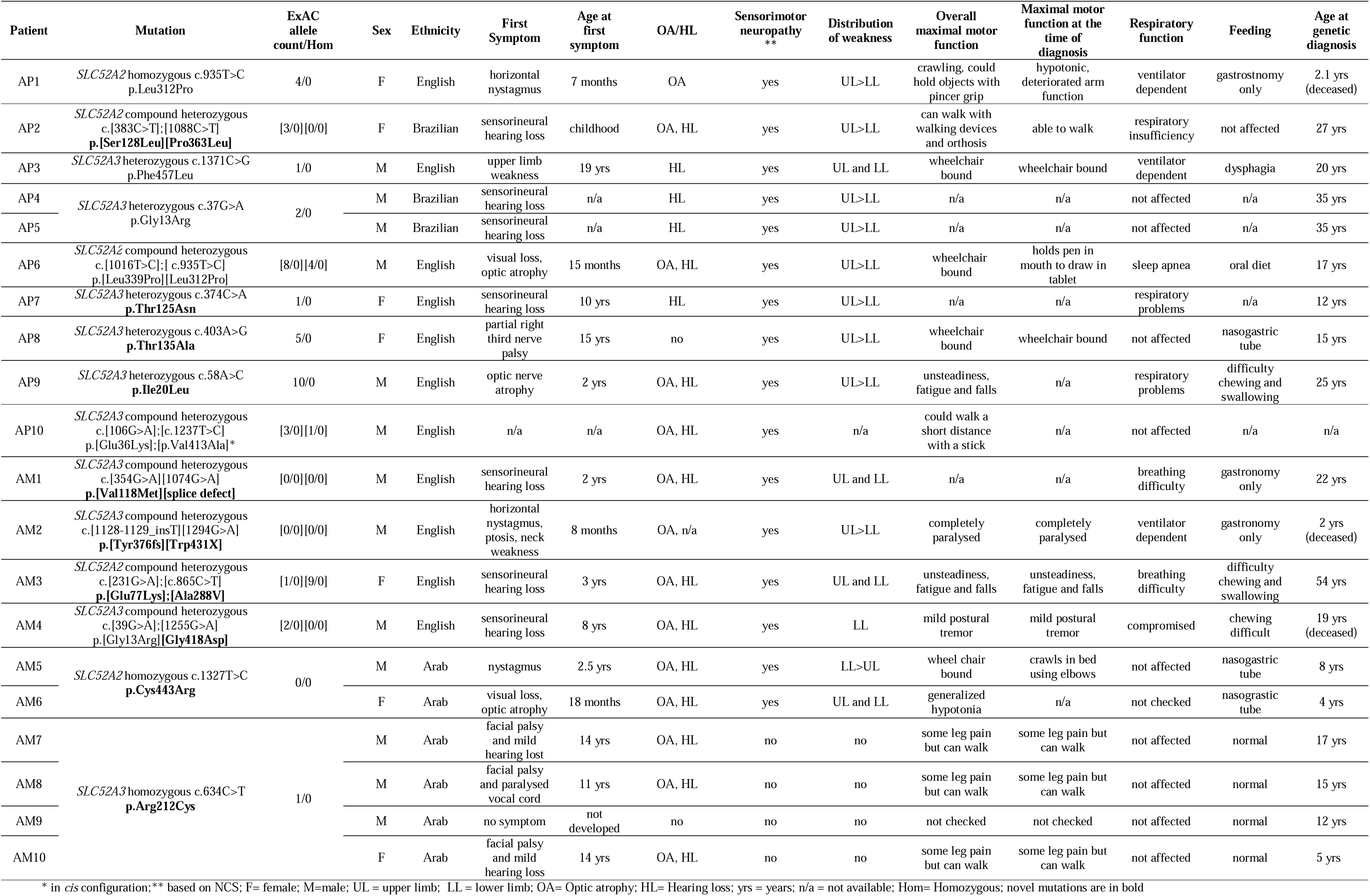
Clinical and genetic features of patients with Brown-Vialetto-Van Laere syndrome caused by mutations in *SLC52A2* and *SLC52A3* at the time of diagnosis. Novel mutations are in bold.

Of the twenty-two mutations identified, eight were found in the *SLC52A2* locus and 14 in *SLC52A3,* of which five *SLC52A2* and nine *SLC52A3* variants were novel (Table 1). In contrast, no *SLC52A1* mutations were observed. None of the variants were found in the homozygous state in the ExAC database of over 100,000 controls, suggesting pathogenicity (Table 1). Consistent with this hypothesis, all except one mutation (p.Arg212Cys in *SLC52A3)* were predicted to be at least probably damaging by SIFT and Polyphen-2 algorithms (scores ranging between 0.55 and 1). The mutations reside in transmembrane helices and in the intracellular and extracellular loops. Although three homozygous and seven compound heterozygous mutations were identified, five mutations (all in *SLC52A3)* were identified on only one allele. These heterozygous individuals did not differ substantially in phenotype including age of presentation from the rest of the cohort of mutation-positive cases.

There was no correlation between the nature of pathogenic variant and phenotype severity, although in the case of patient AM2, nonsense mutations on both alleles resulted in a truncated protein, and this genotype correlated with rapid progression of symptoms and death at 2 years of age.

### Neuropathological analysis of BVVLS patients

To characterise in detail the neuropathological symptoms of BVVLS, we undertook a comprehensive pathological examination of two patients carrying compound heterozygous *SLC52A3* mutations (AM2 and AM4; Table 1). These patients represent two ends of the spectrum of severity of BVVLS.

Patient AM2 had a normal birth at term. His motor skills were mildly delayed and he never acquired the ability to roll over completely front to back. He achieved the ability to sit with minimal support at age 7 months. From about 8 months of age he began to exhibit more clear signs of the condition such as ptosis and neck weakness. He was admitted at the age of 9 months for investigations but no diagnosis was made at that time. His condition quickly progressed to include respiratory muscle weakness, and ventilator dependence at the age of 1 year. He also developed severe weakness in his shoulder girdle areas and proximal upper limbs. Weakness subsequently developed in forearm muscles, distal lower limb muscles and thighs, trunk and face, and progressed to the point that he could only weakly move his eyelids and had very limited sideways movements of his eyes. Nerve conduction studies (NCV) and electromyography showed a severe neuropathy. He died at two years of age of respiratory failure.

The cerebral cortex, hippocampus and cerebral white matter were unremarkable. The deep grey nuclei were not available for assessment. The neuronal density in the substantia nigra was normal for the patient’s age, but the 3^rd^ and 4^th^ cranial nerve nuclei showed severe neuronal loss, gliosis and microglial activation (Supplementary Fig. 1). In the pons there were two symmetrical sharply demarcated lesions surrounding both 5^th^ cranial nerves (Fig. 1A, A1, A2 – D, D1, D2). In these lesions we observed prominent neovascularisation, dense infiltration of macrophages and widespread myelin loss, with relative preservation of axons. The 5^th^ cranial nerve was vacuolated and its nucleus showed severe neuronal loss and gliosis (Supplementary Fig. 1). In the medial lemniscus at the level of the midbrain and pons there was prominent gliosis, microglial activation and vacuolation of the neuropil, which was also seen in the central tegmental tract. In the medulla, the 9^th^, 10^th^ and 12^th^ cranial nerve nuclei showed severe neuronal depletion and gliosis with pale corresponding nerve tracts. The neuronal loss in the 8^th^ cranial nerve was moderate (Supplementary Fig. 1). The medial lemniscus, spinocerebellar tract and medullary reticular formation were all gliotic. Inferior olivary nuclei, in particular the dorsal and ventral parts, showed severe neuronal loss and gliosis. In the cerebellum, there was no significant cortical atrophy (Supplementary Fig. 2B-B1), but the cerebellar white matter showed widespread vacuolation. In the middle cerebellar peduncle there was a lesion with similar morphology to those seen in the pons and medulla. In the spinal cord, there was moderate neuronal loss in the anterior horns and moderately severe atrophy of spinothalamic tracts, spinocerebellar tracts, gracile fasciculus and nucleus in the medulla, and to a lesser extent cuneate fasciculus and nucleus in the medulla (Supplementary Fig. 1). There was severe fibre loss and macrophage activation in the anterior spinal nerve roots, whilst the poster spinal nerve roots were densely populated by myelinated fibres with little macrophage activation (Fig. 1E, E1, E2 – H, H1, H2).

**Figure 1.**
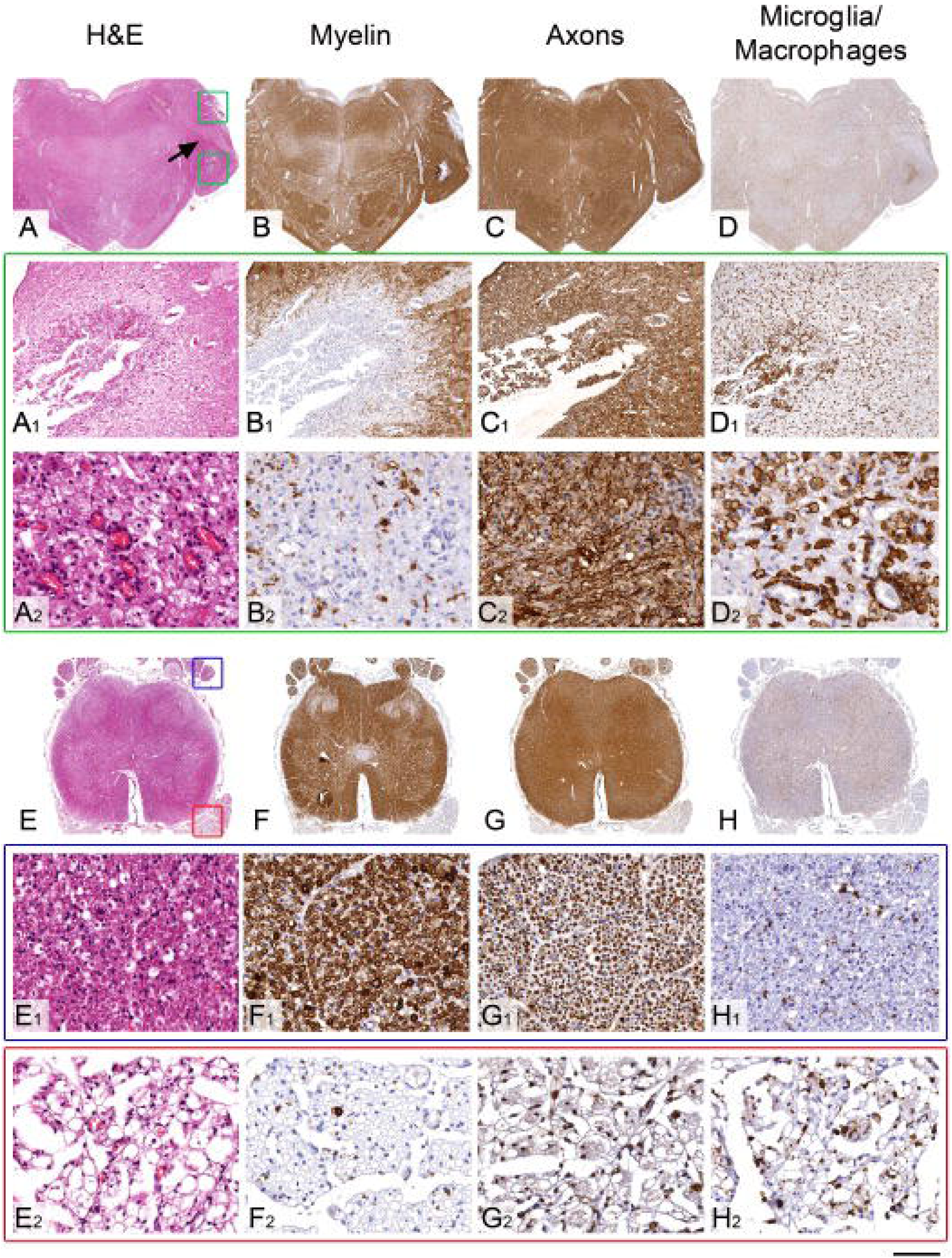
Pathological features of the symmetrical brain stem lesions and the comparison of the spinal cord nerve root involvement in the patient AM2. (A, B, C and D) Low power views of the pons and (A_1_, B_1_, C_1_, D_1_, A_2_, B_2_, C_2_ and D_2_) high power views of the pontine lesion indicated with green square boxes in A. (A, A_1_ and A_2_) The Haematoxylin and Eosin (H&E) stained section demonstrates sharply demarcated lesion surrounding the 5^th^ cranial nerve (black arrow). High power view (A_2_) reveals frequent small calibre blood vessels and foamy macrophages within the lesion. (B, B_1_ and B_2_) Immunostaining for myelin with SMI94 antibody shows a near complete absence of myelin, whilst the axons (C, C_1_ and C_2_), demonstrated with SMI31 antibody, are well preserved within the lesion. (D, D_1_ and D_2_) Immunostaining for macrophages with CD68 antibody confirm a dense infiltrate of macrophage in the centre of the lesion. (E, F, G and H) Low power views of transverse sections of the sacral spinal cord. The posterior nerve roots are indicated with blue square box in E and on high power views in E_1_, F_1_, G_1_ and H_1_. The anterior nerve roots are indicated with red square box in E and on high power views in E_2_, F_2_, G_2_ and H_2_. (E_1_, F_1_, and G_1_) The posterior nerve roots are densely populated with myelinated fibres with (H_1_) minimal macrophage activity. (E_2_, F_2_, and G_2_) In the anterior nerve roots there is a severe loss of myelinated fibres and (H_2_) widespread infiltration of macrophages. Scale bar: 4 mm in A-D; 1.7 mm in E-H; 350 μm in A_1_- D_1_; 70 μm in A_2_- D_2_; 140 μm in E_1_- H_1_ and E_2_- H_2_.

Patient AM4 also had a normal birth at term. He had hearing problems from the age of 8 years and was diagnosed with sensorineural hearing loss at the age of 11. He developed optic atrophy and difficulty walking at the age of 16 years. At age 17 years he presented with dysarthria and subsequently developed swallowing difficulties. Electromyography showed widespread denervation and NCS studies were consistent with an axonal motor-neuropathy. He died at 19 years of respiratory insufficiency.

The cerebral neocortex, hippocampus, amygdala, caudate nucleus putamen, globus pallidus, thalamus and cerebral deep white matter showed no apparent abnormality. Mild gliosis was evident in the dorsal part of the optic tract. In the midbrain, the substantia nigra was densely populated by lightly pigmented neurons in keeping with the patient’s age. The corticospinal tracts were unremarkable in the cerebral peduncles of the midbrain. The 3^rd^ cranial nerve nucleus was not available for the assessment and the 4^th^ cranial nerve nuclei showed a mild degree of neuronal loss and accompanying mild gliosis (Supplementary Fig. 3). In the pons there was prominent neuronal loss in the loci coerulei with free pigment deposits in the neuropil and gliosis (Supplementary Fig. 3). The neuronal density in the pontine basal nuclei and the fibre density in the transverse and corticospinal tracts were unremarkable. The superior cerebellar peduncles showed mild gliosis and vacuolation. The 5^th^ and 7^th^ cranial nerve nuclei showed severe neuronal loss (Supplementary Fig. 3 and Supplementary Fig. 4A, C). In the 6^th^ cranial nerve nucleus the gliosis was mild and the neuronal density was relatively well-preserved (Supplementary Fig. 3 and Supplementary Fig.4B). In the medulla (Supplementary Fig. 3 and Supplementary Fig. 4D-F), the nuclei of the 8^th^ cranial nerve showed severe neuronal loss and prominent gliosis and the nuclei of the 10^th^ and 12^th^ cranial nerves and nucleus ambiguus showed moderately severe neuronal loss and gliosis. The nerve tracts of the available 5^th^, 10^th^ and 12^th^ cranial nerves (Supplementary Fig. 4: insets in A, E and F) showed reduced density of the myelinated fibres. The gracile nucleus showed severe, while the cuneate nucleus showed mild neuronal loss and gliosis (Supplementary Fig. 3). There was a severe pallor, gliosis and prominent microglial/macrophage activation in the medial lemniscus (Supplementary Fig. 4J), spinothalamic tracts and inferior cerebellar peduncles (Supplementary Fig. 4K). The corticospinal tracts in the pyramids (Supplementary Fig. 4L) were densely populated by myelinated fibres and the inferior olivary nuclei showed only mild patchy gliosis (Supplementary Fig. 3). In the cerebellar cortex there was widespread Purkinje cell loss and Bergmann gliosis and some degree of neuronal loss in the granulare cell layer (Supplementary Fig. 2A-A1 and Supplementary Fig. 4G). The cerebellar nuclei - fastigii, globosus and emboliformis - showed severe neuronal loss and gliosis bilaterally, while the dentate nuclei were mildly gliotic with less conspicuous neuronal loss (Supplementary Fig. 4H-I). In the ventral aspect of the upper spinal cord at the junction with the lower medulla there were bilateral symmetric lesions involving the anterior horns, lateral reticular nucleus, supraspinal nucleus, spinothalamic tracts and medial longitudinal fasciculus (Fig. 2A-E). In these lesions there was neovascularization and dense infiltration of macrophages with a near complete loss of myelin, whilst neurons and axons were relatively preserved (Fig. 2A1-E1). Below these lesions the cervical cord showed a severe atrophy of the neurons in the anterior horns and to a lesser extent posterior horns (Supplementary Fig. 3). There was a severe, slightly asymmetrical atrophy of the uncrossed anterior corticospinal tracts, whilst the lateral corticospinal tracts were densely populated by myelinated fibres with only mild vacuolation (Supplementary Fig. 3 and Supplementary Fig. 4L1). Severe symmetrical atrophy with prominent pallor, macrophage infiltration and vacuolation of the anterior and posterior spinocerebellar tracts (Supplementary Fig. 4 K1), spinothalamic tracts and gracile and to a lesser extent cuneate fasciculi (Supplementary Fig. 3) was also observed. Whilst the posterior nerve roots were densely populated by myelinated fibres with only mild macrophage activation, in the anterior nerve roots there was moderately severe loss of myelinated fibres and prominent infiltration of macrophages (Fig. 2F, F1, F2 – J, J1, J2).

**Figure 2.**
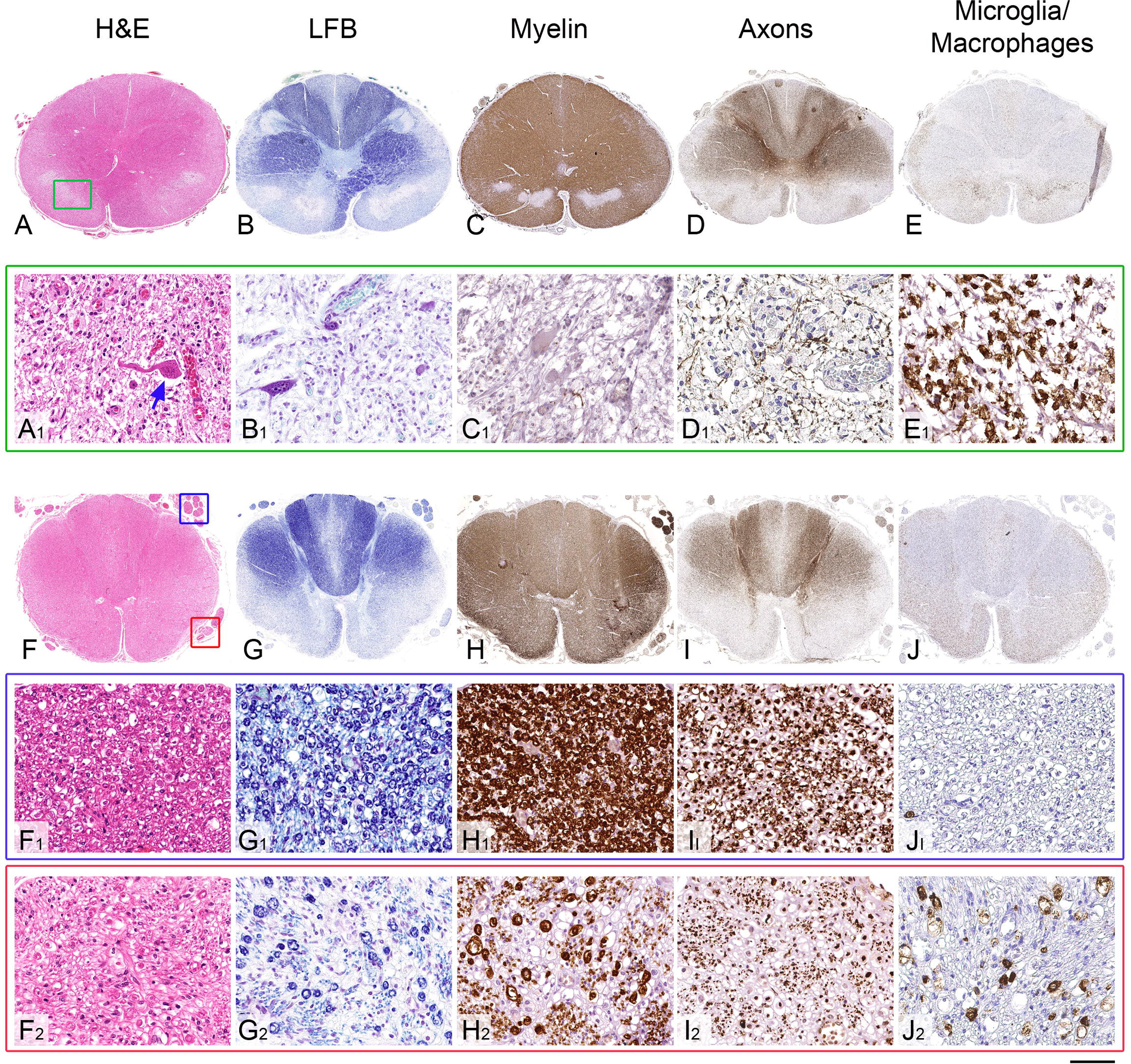
Pathological features of the symmetrical brain stem and spinal cord lesions and the comparison of the spinal cord nerve root involvement in the patient AM4. (A, B, C, D and E) Low power views of the upper cervical cord and (A_1_, B_1_, C_1_, D_1_ and E_1_) high power views of the lesion indicated with green square box in A. (A and A_1_) The H&E stained section demonstrates bilateral symmetrical sharply demarcated lesions (indicated with a green square box on one side) in the anterior part of the upper cervical cord. High power view of the lesion (A_1_) reveals intact neurones (blue arrow) within a dense infiltrate of foamy macrophages and increased numbers of small calibre blood vessels. (B and B_1_) Staining for myelin with luxol fast blue (LFB) special stain and (C and C_1_) immunostaining with SMI94 antibody highlights the preservation of the neurones and shows complete absence of myelin, whilst the axons (D and D_1_), demonstrated with SMI31 antibody, are better preserved within the lesion. (E and E_1_) Dense infiltrates of macrophages within the bilateral lesions are confirmed with CD68 immunohistochemistry. (F, G, H, I and J) Low power views of transverse section of the thoracic spinal cord. The posterior nerve roots are indicated with blue square box in F and on high power views in F_1_, G_1_, H_1_, I_1_ and J_1_. The anterior nerve roots are indicated with red square box in F and on high power views in F_2_, G_2_, H_2_, I_2_ and J_2_. (F_1_, G_1_, H_1_ and I_1_) The posterior nerve roots are densely populated with myelinated fibres with (J_1_) minimal macrophage activity. (F_2_, G_2_, H_2_ and I_2_) In the anterior nerve roots there is a moderately severe loss of myelinated fibres and (J2) prominent macrophage activation. Scale bar: 2.5 mm in A-E and F-J; 80 μm in A_1_- E_1;_ F_1_- J_1_ and F_2_- J_2_.

Neither in the case AM2 nor AM4 were any amyloid-β, hyper-phosphorylated tau, α-synuclein, TDP-43, p62 or ubiquitin positive inclusions observed. In both cases the inflammatory reaction was restricted to microglial activation and macrophage infiltrates with no significant lymphocytic inflammation.

### Fibroblasts biochemical studies

Mitochondrial dysfunction has long been documented in neurodegenerative diseases (Palomo and Manfredi, 2015). For example, SOD1 mutations in familial ALS have been shown to lead to abnormalities in mitochondrial morphology, both in biopsies and post-mortem tissues of human patients (Sasaki and Iwata, 2007; Sasaki, 2010) and in cellular and mouse models of the disease (Magrane et al., 2009; Magrane et al., 2014; Palomo and Manfredi, 2015; Vinsant et al., 2013). However, whether mitochondrial function is perturbed by BVVLS-linked mutations has yet to be examined. We hypothesized that lower levels of intracellular riboflavin as a result of mutated RFVTs would lead to reduced levels of FMN and FAD, which in turn would lead to impairments at the level of the ETC complex I and complex II. Using fibroblasts derived from BVVLS patients with RFVT mutations and healthy age-matched controls, we found a significant reduction in the intracellular levels of FMN and FAD in patient fibroblasts when grown in low extracellular riboflavin conditions (Fig. 3A, B). Levels of intracellular riboflavin in patient fibroblasts frequently fell below the threshold of detection under these conditions (but not in control fibroblasts; data not shown), consistent with defective riboflavin transport. Furthermore, we observed a significant reduction in ETC complex I and complex II activity in patient fibroblasts compared to controls (Fig. 3C, D).

**Figure 3.**
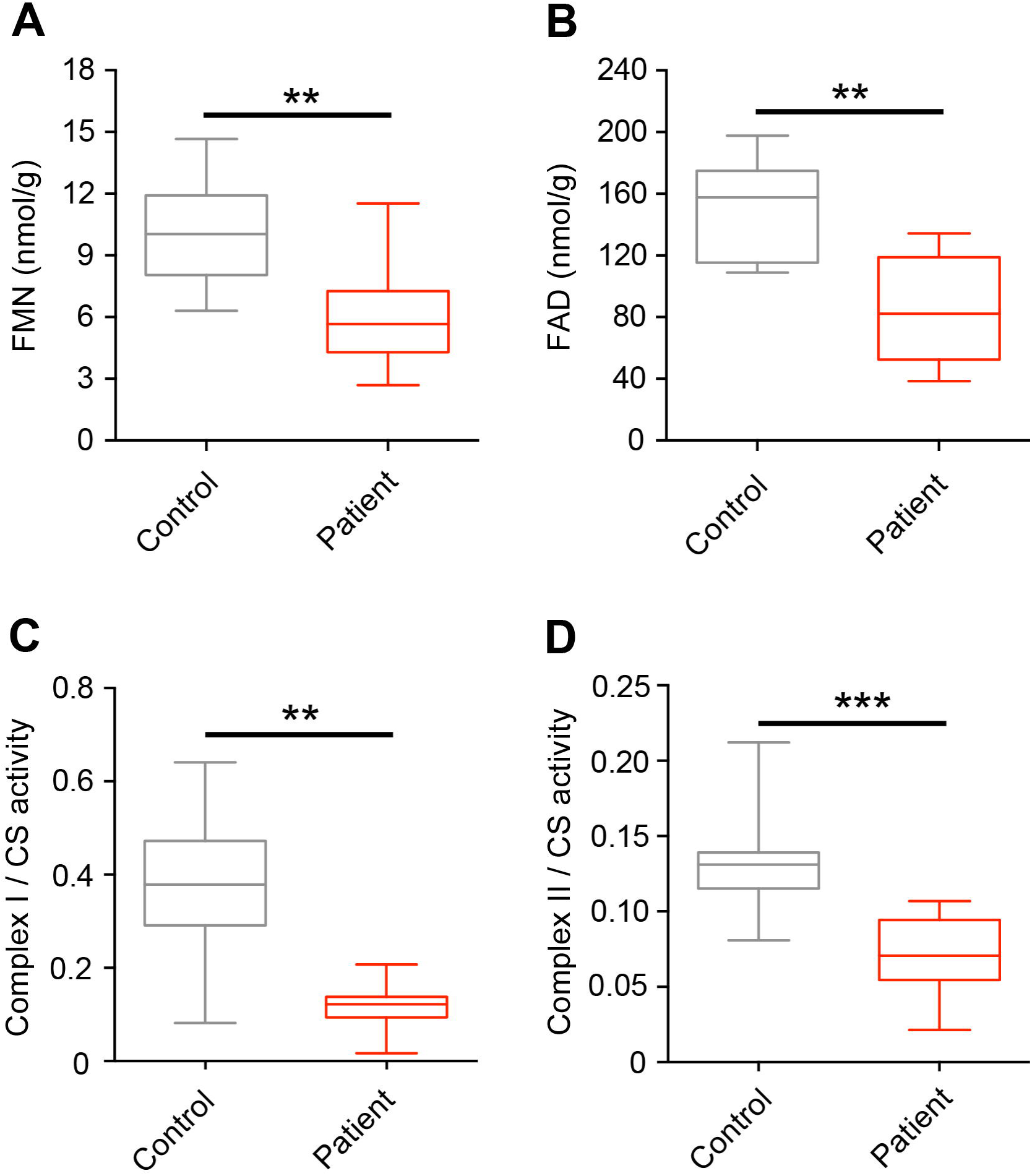
Reduced mitochondrial activity in BVVLS patient fibroblasts. (A-B) Intracellular FMN (A) and FAD (B) levels in control and patient fibroblasts. (C-D) Complex I (C) and complex II (D) activity in control and patients’ fibroblasts. Complex activities are expressed as a ratio to citrate synthase activity. Data are presented as box plots illustrating 80% of the data distribution. 10^th^, 25^th^, Median, 75^th^ and 90^th^ percentiles are shown in these and all subsequent box plots. **p < 0.005, ***p < 0.0005, Mann-Whitney U-test. Data were generated from a minimum of three independent experiments. n = 3 and n = 5 for control and patient fibroblasts respectively.

### Generation of a novel *Drosophila* model of BVVLS

We next sought to confirm the link between RFVT dysfunction and reduced mitochondrial activity *in vivo.* Previous work has utilised knock-out of the mouse *SLC52A3* orthologue to model BVVLS (Intoh et al., 2016; Yoshimatsu et al., 2016). However, *SLC52A3* knock-out mice are lethal within 48 h of birth (Yoshimatsu et al., 2016), precluding phenotypic analysis at later developmental stages. To generate a new *in vivo* model of BVVLS, we turned to the fruit fly, *Drosophila melanogaster*. Comparative BLAST analysis revealed a single *Drosophila* homologue of SLC52A3/hRFVT3, the previously uncharacterised gene *cg11576* (E-value: 2.74e^-74^; next closest homologue, E-value: 2.95). As described below, we name the *cg11576* gene *drift* (*<u>D</u>rosophila* <u>r</u>ibo<u>f</u>lavin <u>t</u>ransporter). The DRIFT amino-acid sequence exhibits 36.9% identity and 53.1% similarity with hRFVT3, and the L1 loop and GXXXG motifs characteristic of RFVTs are also conserved (Fig. 4A and Supplementary Fig. 5A) (Russ and Engelman, 2000; Zhang et al., 2010). The L1 loop is a region of the protein shown to recognize riboflavin through both hydrogen bonds and van der Waals interactions (Zhang et al., 2010), while the GXXXG motif is required for dimerization (Russ and Engelman, 2000). DRIFT is also highly homologous to the hRFVT3 paralogues hRFVT1 and hRFVT2 (Fig. 4A). Furthermore, 10/19 and 11/25 residues in hRFVT2 and hRFVT3 respectively that are altered by BVVLS-linked mutations are either identical or functionally similar in DRIFT. Mammalian RFVTs exhibit wide domains of expression, including the nervous system, intestine, testes, and placenta (Yonezawa and Inui, 2013). Similarly, *drift* is transcribed in several adult *Drosophila* tissues, including the head, gut, abdomen and thorax (Supplementary Fig. 5B). This broad expression pattern is in agreement with RNAseq data from the *Drosophila* ModEncode Project (Graveley et al., 2011).

**Figure 4.**
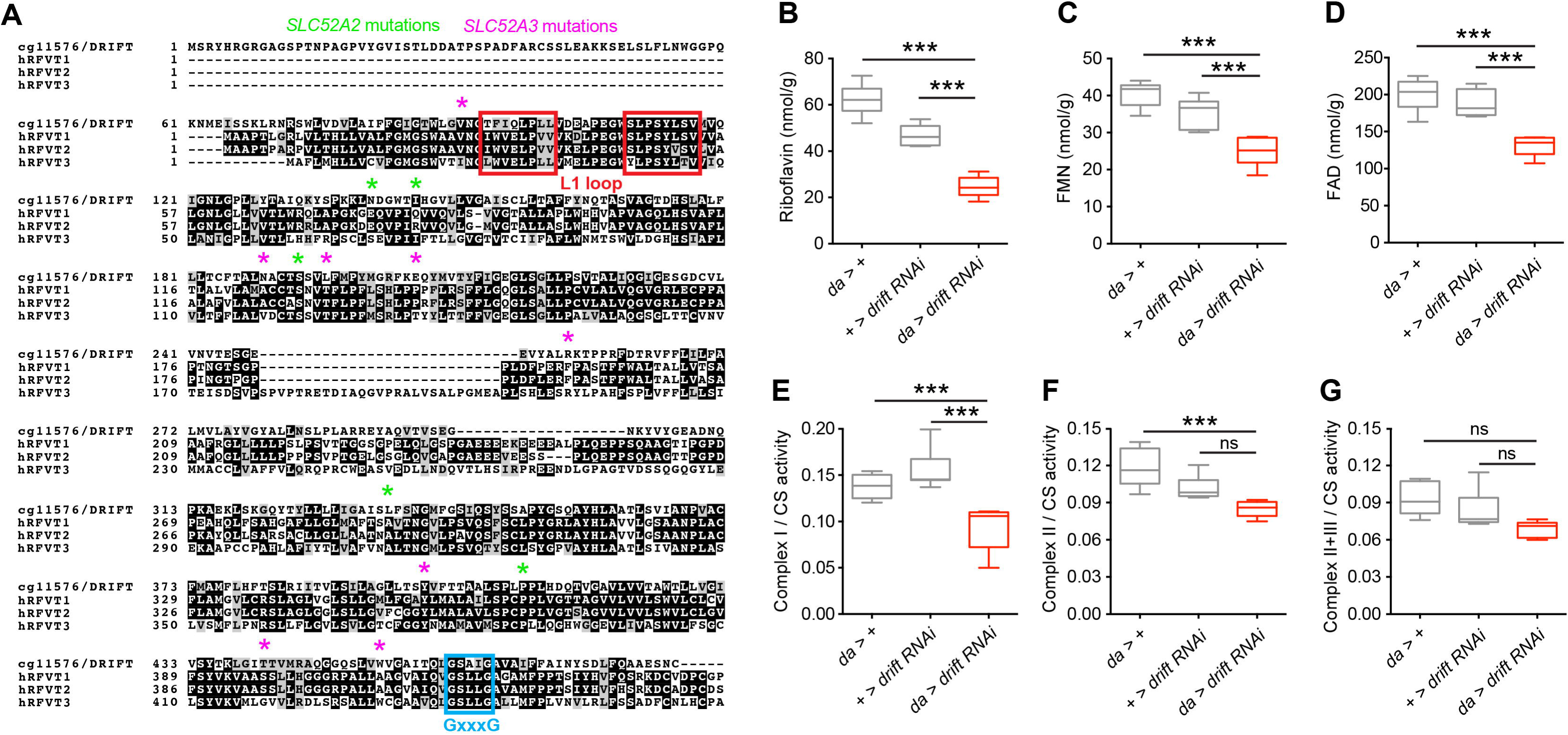
Knockdown of the *Drosophila* RFVT homologue *drift* results in reduced mitochondrial activity. (A) Amino acid homology between *Drosophila* DRIFT and human hRFVTl, hRFVT2 and hRFVT3. Novel mutations found in this study are represented in green for *SLC52A2* (hRFVT2) and pink for *SLC52A3* (hRFVT3). Black: identical amino acid, Grey: functionally similar amino acids. The L1 loops (red) and GXXXG motifs (blue) characteristic of RFVTs are shown. Conservation among species of the amino acid residues was determined using ClustalW2 software for multiple sequence alignment and plotted with BOXSHADE. B-D) Riboflavin (B), FMN (C) and FAD (D) levels in *drift* knockdown flies and controls normalised to total protein levels. (E-G) Complex I (E), complex II (F) and complex II-III (G) activity in *drift* knockdown flies and controls. Complex activities are normalised to citrate synthase (CS) activity. ***p < 0.0005, ns - p > 0.05, one-way ANOVA with Dunnett’s post-hoc test. Data were generated from a minimum of three independent experiments. n = 10 for each genotype. Individual measurements were performed in duplicates.

To examine the biochemical and phenotypic consequences of loss of DRIFT, we disrupted *drift* expression using either P-element insertions or transgenic RNA interference (RNAi). Insertion of a MiMIC P-element insertion (*cg11576*^MI04904-GFSTF.2^) within intron 1 of *drift* resulted in viable heterozygote adults but homozygote lethality prior to the L3 larval stage. Furthermore, global knockdown of *drift* using two independent transgenic RNAi lines (7578 and HMC04813) driven via a strong global promoter (*actin*-Gal4) also resulted in either complete or highly penetrant lethality prior to the adult stage. Thus, similarly to *SLC52A3* (Intoh et al., 2016; Yonezawa and Inui, 2013; Yoshimatsu et al., 2016), and consistent with a fundamental requirement for riboflavin transport in metazoans (Biswas et al., 2013), DRIFT is likely to be essential in *Drosophila.*

Using a distinct global driver (*daughterless*-Gal4; da-Gal4) in concert with the 7578 RNAi line, we achieved robust *drift* knockdown as determined by semi-quantitative RT-PCR (Supplementary Fig. 5C). However, this knockdown did not result in early lethality (suggesting stronger RNAi expression by *actin-* relative to da-Gal4), facilitating analysis of *drift* knockdown flies at later developmental stages. Using whole-body tissue from *drift* knockdown adults and associated control lines (heterozygotes for *da-* Gal4 and the *drift* RNAi transgene), we found that *drift* knockdown resulted in a substantial reduction of *in vivo* riboflavin levels as well as the riboflavin metabolites FMN and FAD (Fig. 4B-D). These results, combined with the high homology of DRIFT to hRFVT1-3, strongly suggest that DRIFT is a *bona fide* riboflavin transporter. We next asked whether *drift* knockdown resulted in reduced mitochondrial activity. Similarly to BVVLS patient fibroblasts, ETC complex I activity was profoundly reduced by *drift* knockdown (Fig. 4E), while ETC complex II and II-III activity exhibit a trend towards lower levels, albeit non-significant (Fig. 4F, G). Thus, in *Drosophila,* complex I activity appears particularly sensitive to DRIFT expression.

### *drift* knockdown results in reduced locomotion and lifespan in *Drosophila*

The viability resulting from *drift* knockdown via da-Gal4 allowed us to assess whether RFVT knockdown impacts post-embryonic organismal phenotypes in a manner consistent with BVVLS pathology (Foley et al., 2014; Manole et al., 2014). We found that *drift* knockdown resulted in profound locomotor defects in both larval and adult *Drosophila. drift* knockdown 3^rd^ instar larvae exhibit significantly reduced locomotion, as measured by the number of grid crosses per minute on an agar plate (Supplementary Fig. 5D). We also used the *Drosophila* activity monitor (DAM) system to perform automated recordings of adult locomotion, measured as the number of infra-red beam breaks across a 24 h day/night cycle. Under 12 h light: 12 h dark conditions, control 1-2 day old adult female *Drosophila* exhibit peaks of activity at dawn and dusk, and relative quiescence during the afternoon and night (Fig. 5A, B). In contrast, peak activity and total beam breaks in *drift* knockdown adults were substantially reduced (Fig. 5C, D). Furthermore, *drift* knockdown resulted in greatly reduced lifespan, with 99% mortality within four days post-eclosion (Fig. 5E). These phenotypes mimic motor problems and early mortality observed in BVVLS patients, suggesting a conserved link between RFVT dysfunction, locomotor strength and lifespan.

**Figure 5.**
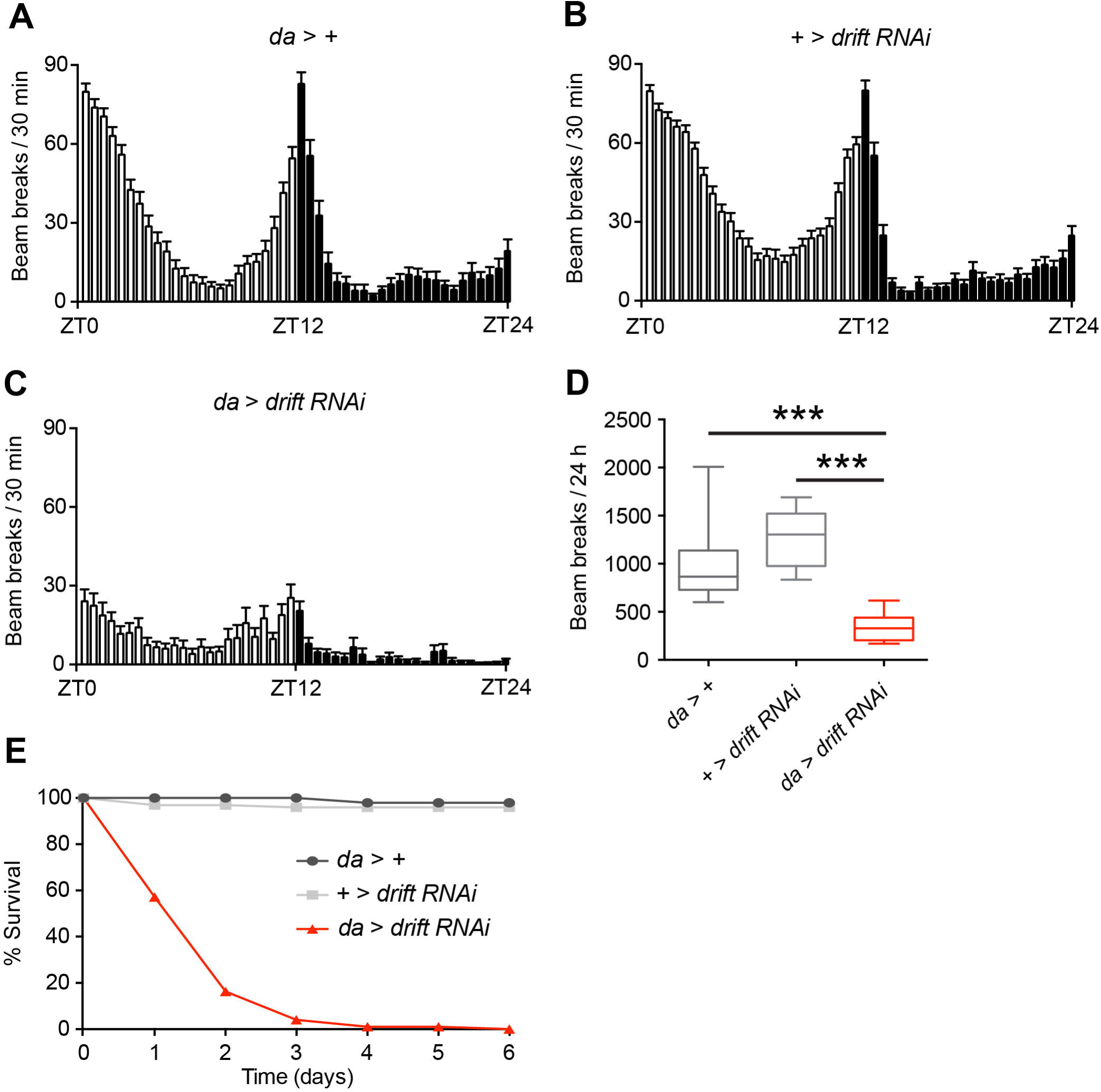
*drift* knockdown reduces locomotor activity and lifespan in adult *Drosophila*. (A-C) Locomotor activity over 24 h of driver alone (A) and drift RNAi alone (B) controls, and *drift* knockdown flies (C). Data was derived from adult females. Mean values for each time point are presented; error bars indicate standard error of the mean (SEM). (D) Box plots illustrating total activity over 24 h for each genotype. ***p < 0.0005, Kruskal-Wallis test with Dunn’s post-hoc test. Data were generated at least three independent experiments. n = 30 for each control and n = 16 for the *drift* knockdown flies. (E) Percentage survival of *drift* knockdown flies and controls on normal food. n = 98 for each control and n = 99 for the *drift* knockdown flies.

### An esterified riboflavin derivative partially rescues *drift* knockdown phenotypes

Since BVVLS pathology can be partially ameliorated by riboflavin treatment, we asked whether locomotor defects in *drift* knockdown flies could be rescued by supplementing *Drosophila* culture medium with riboflavin (0.1 mg/ml). However, we found no enhancement of locomotor activity following riboflavin supplementation (Supplementary Fig. 5E). Riboflavin is a water-soluble vitamin that is easily excreted, leading to low bioavailability and short half-life. Furthermore, since RFVT expression is very low in *drift* knockdown flies (Supplementary Fig. 5C), riboflavin in the *Drosophila* haemolymph may fail to be transported into relevant cell-types. We sought to circumvent these issues using an esterified derivative of riboflavin (riboflavin-5’-lauric acid monoester; RLAM; 0.1 mg/ml) that could act as a pro-drug, likely diffuse into the intracellular space independently of RFVT function and be cleaved by esterases to release active riboflavin (Fig. 6A). As predicted, food supplementation with RLAM dramatically increased complex I activity (~ 3-fold), and critically, resulted in heightened total locomotion (Fig. 6B, C), increased peak levels of daily activity (Fig. 6D, E), and a partial extension of lifespan (Fig. 6F) in *drift* knockdown flies. We speculate that RLAM may represent a more efficient treatment method for BVVLS patients since cellular uptake of RLAM may still robustly occur in the absence of functional endogenous RFVTs.

**Figure 6.**
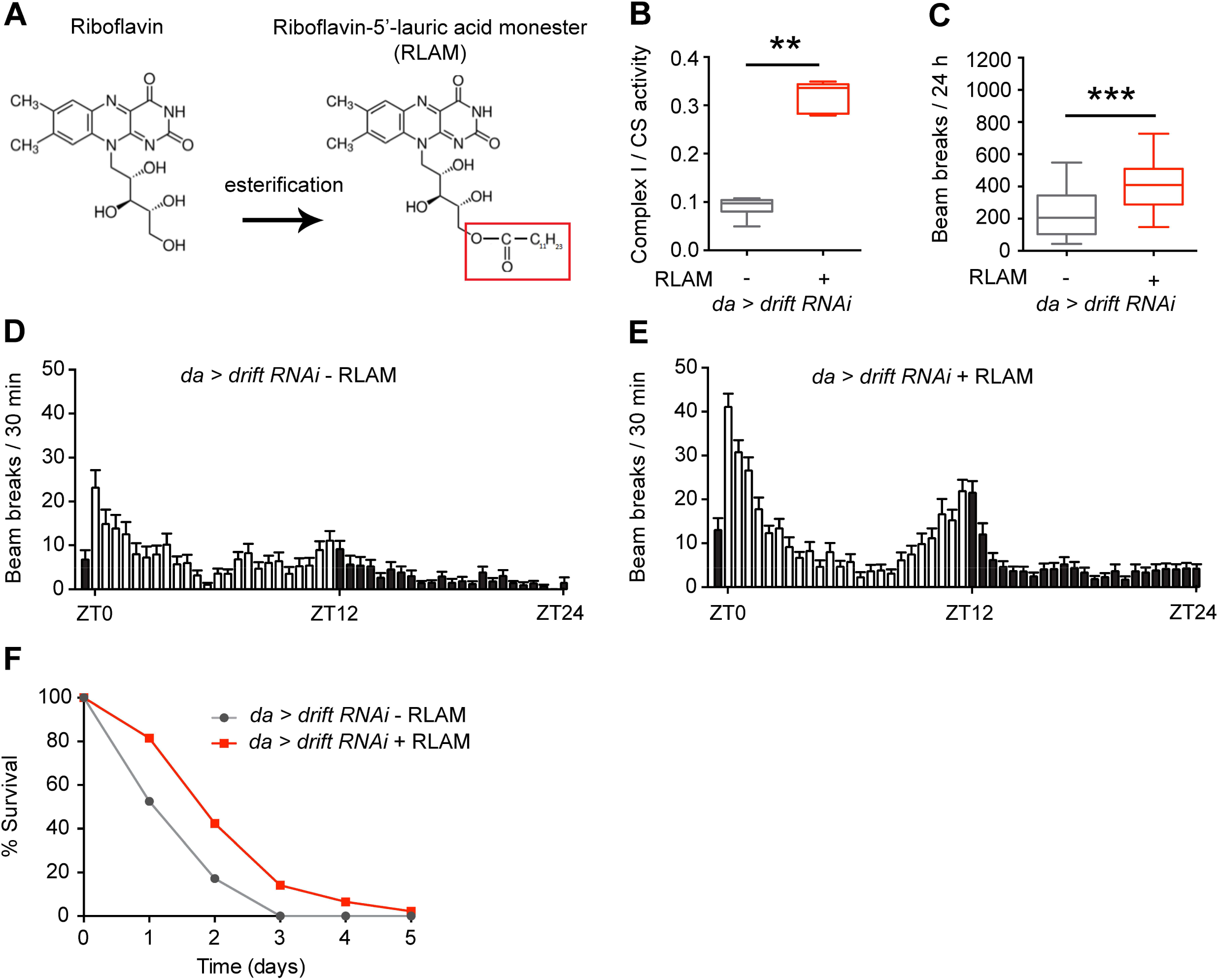
A riboflavin ester partially rescues the *drift* knockdown phenotypes. (A) Chemical structure of riboflavin and its ester (RLAM). (B) Complex I activity in *drift* knockdown flies grown on normal and RLAM-supplemented food. Complex activity is expressed as a ratio to citrate synthase (CS) activity. Data were generated at least three independent experiments. n = 10 for each genotype. Individual measurements were done in duplicates. (C) Total locomotor activity over 24 h in *drift* knockdown grown on normal and RLAM-supplemented food. **p < 0.005, ***p < 0.0005, Mann-Whitney U-test. Data were generated at least three independent experiments. n = 37 and n = 52 *drift* knockdown fed normal and RLAM-supplemented conditions respectively. (D-E) Mean locomotor activity over 24 hours of the *drift* knockdown flies grown on normal food (D) and RLAM-supplemented food (E). Values are presented as a mean and SEM. (F) Percentage survival of *drift* knockdown flies grown on normal and RLAM-supplemented food. n = 99 and n = 92 *drift* knockdown fed normal and RLAM-supplemented conditions respectively.

## Discussion

We expanded the spectrum of genetic defects in RFVTs identifying *SLC52A2* and *SLC52A3* mutations in 15% of cases. It was previously noted that the distinct phenotype of upper limb and axial weakness, hearing loss and optic atrophy could be attributed only to patients harbouring *SLC52A2* mutations (Foley et al., 2014). However, in our cohort of 20 patients, mutations in both *SLC52A2* and *SLC52A3* resulted in similar phenotypes. *SLC52A3* mutations were more frequent than mutations in *SLC52A2* in our cohort. This is in agreement with previous reports where BVVLS-linked mutations were predominantly found in *SLC52A3* (Bosch et al., 2011; Manole and Houlden, 2015). The high proportion of patients who were found to be negative for mutations in the known RFVTs, indicates that novel genetic causes are yet to be found.

The neuropathology of BVVLS has not been fully characterised in the past (Malafronte et al., 2013). In keeping with the spectrum of clinical findings, severe depletion of neurons in multiple cranial nerve nuclei, anterior horns of the spinal cord, cerebellar nuclei, Purkinje cells in the cerebellum, and degeneration of optic pathway, solitary tract, and spinocerebellar and pyramidal tracts was seen along with an axonal neuropathy in the sural nerve (Foley et al., 2014). Here we provide detailed neuropathological assessment of the brains and spinal cords in two patients with genetically confirmed mutations in the *SLC52A3* gene, who fell at the two extremes of age of presentation of BVVLS. There was variably and severe neuronal loss and gliosis in the brain stem cranial nerve nuclei and anterior horns of the spinal cord with accompanying nerve root atrophy which reflects the spectrum of clinical symptomatology. Whilst no correlation between the mutation type and severity of clinical phenotypes was evident in the case series presented here, it is possible that different mutations directly influence the length of the disease and degree of atrophy of specific brain structures. For example, the rapid deterioration and demise of patient AM2 due to formation of truncated protein originating from nonsense mutations on both alleles may explain less prominent atrophy of cerebellar Purkinje cells when compared with patient AM4, the atrophy of which may require longer duration of the disease (Supplementary Fig. 2). In both patients there were symmetrical lesions in the brain stem characterised by prominent neovascularisation, dense infiltration of macrophages, loss of myelin and relative preservation of neurons and axons. Although the anatomical distribution of the symmetrical lesions differed in both cases (Supplementary Figs. 1 and 3), the morphology of the lesions was identical in both cases and was similar to the pathology seen in mitochondrial encephalopathies (Filosto et al., 2007; Tanji et al., 2001). To the best of our knowledge such lesions have not been documented in any of the previous published cases of BVVL clinical syndrome and link with the mitochondrial abnormalities in flies and patient fibroblasts.

Although fibroblasts are non-neural cells and not particularly vulnerable as a tissue, metabolic and mitochondrial abnormalities are commonly studied in this cell-type (Distelmaier et al., 2009). We identified clear deficiencies in the activities of ETC complex I and complex II in patient-derived fibroblasts relative to fibroblasts from healthy controls. Interestingly, previous results have only shown evidence of marginally decreased ETC complex I activity in muscle cells from some *SLC52A2* patients sensitive to reduced riboflavin uptake (Foley et al., 2014). Mitochondrial activity in different tissues and cell types may thus be differentially sensitive to reduced riboflavin uptake.

Finally, we examined the *in vivo* consequences of knockdown of the single *SLC52A3* homologue *drift* in *Drosophila.* Biochemical analysis of *drift* knockdown tissue is consistent with data derived from patient fibroblasts, showing diminished levels of riboflavin, FMN and FAD, and reduced complex I activity. At the whole-organismal level, there was impairment in locomotion at both the larval and adult stages, reminiscent of the limb weakness and movement impairment of the BVVLS patients. Moreover, *drift* knockdown flies had severely reduced life span, similar to untreated patients (Bosch et al., 2011). It is interesting to note that these lifespan and locomotor defects are rescued by ingestion of an esterified derivative of riboflavin, since this provides further evidence that the phenotypic signatures of *drift* knockdown flies are linked to riboflavin deficiency and consequent downstream metabolic defects. We hypothesize that lack of RLAM during pupation, a critical neurodevelopmental stage during which RLAM-supplemented food will not be consumed, may contribute to the partial nature of the observed RLAM rescue. Nonetheless, RLAM fulfils the criteria for being a treatment targeting energy metabolism, a fundamental mitochondrial process in addition to circumventing the disease-related protein, features generally believed to be of great promise (Foley et al., 2014; Lin and Beal, 2006; Xuan et al., 2013).

It was previously thought that defective mitochondria occur secondary to primary disease mechanism, but current research suggests that mitochondrial dysfunction may play a role in both onset and development of neurodegenerative diseases such as Alzheimer’s, Parkinson’s, Huntington’s and ALS, where decreases in one or more mitochondrial complexes and corresponding oxidative stress have been reported in cell or animal models (Fukui and Moraes, 2007; Golpich et al., 2016; Hoglinger et al., 2003; McInnes, 2013; Menzies et al., 2002; Yamada et al., 2014). Our results from *in vitro* and *in vivo* models of a childhood neuropathy are in line with these findings, highlight a contribution of mitochondrial dysfunction to BVVLS pathology, and suggest future therapeutic strategies based on esterified riboflavin.

## Acknowledgements

The authors would like to thank the participants of the study for their essential help with this work. This study was supported by the Medical Research Council (MRC UK), The Wellcome Trust (equipment and the Synaptopathies strategic award (104033)), Ataxia UK, The BRT and The UK HSP Society and the EU FP7/2007-2013 under grant agreement number 2012-305121 (NEUROMICS). The NIH Undiagnosed Diseases Program (UDP) is supported by the Intramural Research Program of NHGRI. We are also supported by the National Institute for Health Research (NIHR) University College London Hospitals (UCLH) Biomedical Research Centre (BRC). RH is a Wellcome Trust Investigator (109915/Z/15/Z), who receives support from the Medical Research Council (UK) (MR/N025431/1), the European Research Council (309548), the Wellcome Trust Pathfinder Scheme (201064/Z/16/Z) and the Newton Fund (UK/Turkey, MR/N027302/1). PFC is a Wellcome Trust Senior Fellow in Clinical Science (101876/Z/13/Z), and a UK NIHR Senior Investigator, who receives support from the Medical Research Council Mitochondrial Biology Unit (MC_UP_1501/2), the Wellcome Trust Centre for Mitochondrial Research (096919Z/11/Z), the Medical Research Council (UK) Centre for Translational Muscle Disease (G0601943), EU FP7 TIRCON, and the National Institute for Health Research (NIHR) Biomedical Research Centre based at Cambridge University Hospitals NHS Foundation Trust and the University of Cambridge. The views expressed are those of the author(s) and not necessarily those of the NHS, the NIHR or the Department of Health.

---

**Figure S1. Schematic representation of the anatomical distribution and severity of the brain stem and spinal cord pathology in the patient AM2.**

In the schematic figures the severity of grey matter pathology is indicated in orange with the lightest shade corresponding to mild neuronal atrophy and mild gliosis and the darkest shade corresponding to severe atrophy and gliosis. The severity of white matter tract pathology is indicated in green with the lightest shade corresponding to mild pathology and the darkest shade corresponding to a severe myelinated fibre loss. The corresponding transverse brain stem and spinal cord sections are immune-stained for myelin with SMI94 antibody. The red arrowheads in the pons indicate the bilateral symmetrical lesions surrounding both 5^th^ cranial nerves, indicated with a yellow arrow on one side. Also the lesion in the medulla is indicated with a red arrowhead.

**Figure S2. Comparison of the cerebellar atrophy between patient AM4 and AM2**

(A and A_1_) correspond to patient AM4 and (B and B_1_) correspond to patient AM2. (A and A_1_) In the cerebellar cortex from the patient AM4 there is severe Purkinje cell loss with widespread Bergmann gliosis. (B and B_1_) In the patient AM2 the Purkinje cells are well preserved. Note the presence of external granular cell layer (B_1_) in patient AM2, a normal finding for the patient’s young age. Scale bar: 80 μm in A-B and 30 μm in A_1_-B_1_.

**Figure S3. Schematic representation of the anatomical distribution and severity of the brain stem and spinal cord pathology in the patient AM4.**

In the schematic figures the severity of grey matter pathology is indicated in orange with the lightest shade corresponding to mild neuronal atrophy and mild gliosis and the darkest shade corresponding to severe atrophy and gliosis. The severity of white matter tract pathology is indicated in green with the lightest shade corresponding to mild and the darkest shade corresponding to a severe fibre loss. The corresponding transverse brain stem and spinal cord sections are stained with luxol fast blue special stain, where normally myelinated tracts are stained dark blue. The red arrowheads in the anterior aspect of the lower medulla and spinal cord indicate the bilateral symmetrical lesions.

**Figure S4. The spectrum of the atrophy in the cranial nerve nuclei, deep cerebellar nuclei and white matter tracts in the patient AM4.**

(A) In the 5^th^ cranial nerve nucleus there is a moderately severe neuronal loss and gliosis and accompanying myelinated fibre loss in the nerve tract (inset). (B) The 6^th^ cranial nerve nucleus shows only very mild neuronal loss and gliosis. (C) In the 7^th^ and (D) 8^th^ cranial nerve nuclei the neuronal loss and accompanying gliosis is very severe. (E) In the nuclei of the 10^th^ and (F) 12^th^ cranial nerves the neuronal loss and gliosis is moderately severe, but the nerve tracts (inset in E for the 10^th^ nerve and inset in F for the 12^th^ nerve) show marked depletion of myelinated fibres. (G) In the cerebellar cortex there is widespread Purkinje cell loss, Bergmann gliosis and gliosis in the molecular cell layer. (H) The globose nucleus shows a severe neuronal loss and gliosis, whilst (I) the neurons in the dentate nucleus are better preserved and gliosis is mild. (J) The medial lemniscus in the medulla, (J_1_) the gracile fasciculus in the posterior column, (K) inferior cerebellar peduncle in hindbrain and spinocerebellar tract in the spinal cord (K_1_) show severe gliosis and vacuolation of the neuropil on H&E stained sections, and microglial activation on immunohistochemistry (not shown). (L and L_1_) The corticospinal tracts at the level of medulla (L) and spinal cord (L_1_) in contrast is well populated by myelinated fibres with no apparent gliosis. Scale bar: 110 μm in A-F and H-L; 220 μm in G. (A-F): stained with luxol fast blue. (G-I and J, J_1_-L, L_1_): stained with H&E

**Figure S5. Knockdown of the *Drosophila* RFVT homologue *drift.***

(A) Structural conservation of amino acids between DRIFT, hRFVT1, hRFVT2 and hRFVT3. Known mutations previously reported (Manole and Houlden, 2015) are represented in green for *SLC52A2* and pink for *SLC52A3.* (B) Expression of *drift* in head, gut, abdomen and thorax of the adult fly, detected through RT-PCR. The expected size of the bands are ~ 620 bp and ~ 370 bp for *drift* and *rp49* respectively. DNA ladder (left) is in bp. (C) Semi-quantitative RT-PCR illustrating *drift* knockdown by transgenic RNAi expressed using the global driver *daughterless-GAL4 (da). da* > + and + > *drift* RNAi are the controls for the driver and RNAi line respectively while the knockdown is represented by the da > *drift* RNAi. Data were generated from three independent biological samples (1-3), each with two technical replicates. (D) The number of grid breaks per min of wandering 3^rd^ instar larvae.

Data are presented as box plots illustrating 80% of the data distribution. 10^th^, 25^th^, Median, 75^th^ and 90^th^ percentiles are shown. ***p < 0.0005, Kruskal-Wallis test with Dunn’s post-hoc test. n = 30 for each genotype. (E). Total activity of *drift* knockdown adult female *Drosophila* over 24 h fed normal and riboflavin-supplemented conditions. ns – p > 0.05, Mann-Whitney U-test. n = 15 and n =17 for *drift* knockdown fed normal and riboflavin-supplemented conditions respectively.

